# High throughput expression profiling and in situ screening of circular RNAs in tissues

**DOI:** 10.1101/183020

**Authors:** Ammar Zaghlool, Adam Ameur, Chenglin Wu, Jakub Orzechowski Westholm, Adnan Niazi, Manimozhi Manivannan, Kelli Bramlett, Mats Nilsson, Lars Feuk

**Author notes:** Corresponding author, Lars Feuk, Dept. of Immunology Genetics and Pathology, Box 815, BMC B11:4, Uppsala University, 751 08 Uppsala University, Sweden.

## Abstract

Circular RNAs (circRNAs) were recently discovered as a class of widely expressed noncoding RNA and have been implicated in regulation of gene expression. However, the function of the majority of circRNAs remains unknown. Studies of circRNAs have been hampered by a lack of essential approaches for detection, quantification and visualization. We therefore developed a target-enrichment sequencing method suitable for high-throughput screening of circRNAs and their linear counterparts. We also applied padlock probes and in situ sequencing to visualize and determine circRNAs localization in human brain tissue at subcellular levels. We measured circRNA abundance across different human samples and tissues. Our results demonstrate the potential of this RNA class to act as a specific diagnostic marker in blood and serum, by detection of circRNAs from genes exclusively expressed in the brain. The powerful and scalable tools we present will enable studies of circRNA function and facilitate screening of circRNA as diagnostic biomarkers.

## Main text

### Background

Circular RNAs (circRNAs) were recently discovered as a novel class of noncoding RNA [1–3]. It is believed that most circRNAs are produced from precursor mRNA through back-splicing of exons, which mainly occurs at annotated exon boundaries, forming covalently closed single-stranded circular transcripts [4]. Although circRNAs were originally reported more than two decades ago, only few examples were found, and circRNA were mostly considered to be splicing noise [5, 6]. However, with the recent advances in RNA sequencing technologies and newly developed bioinformatics approaches, several reports revealed that circRNA expression is widespread in human tissues and appears to be a general feature in all eukaryotes [1-3, 7-11]. It is estimated that as much as 10% of expressed genes in examined cells and tissues produce circRNAs, often times in a cell or tissue and developmental-stage specific manner [1, 3, 7, 11–14]. Even though circRNA are often expressed at low levels, there are at least 100 genes express circRNAs as the predominant transcript isoform [10].

Although the function of the vast majority of circRNAs is still unknown, several studies demonstrated that circRNAs could function as competing endogenous RNAs or microRNA sponges, exemplified by the characterization of *CDR1as* and *SRY* circRNAs [8, 15]. Moreover, emerging evidence suggests potential roles for circRNAs in important physiological functions, for example, in the nervous system [11, 16–19]. In addition to their role as microRNA sponges, circRNAs are believed to regulate RNA polymerase II transcription, and compete with the canonical splicing machinery [14, 18–20]. Recently, several reports provided in vivo and in vitro evidence in multiple metazoan species, including human, that a subset of circRNAs are translated into proteins, thereby adding a new layer to the complexity of eukaryotic gene and protein expression [21–23].

CircRNAs are extraordinarily enriched in the mammalian brain, particularly in the synapses, and are upregulated during neuronal differentiation [11, 24], and have been demonstrated to be involved in the function of the brain [24]. In addition, circRNAs have been shown to accumulate with age in Drosophila [9] and mouse brains [25]. Recent data suggest that circRNAs may play crucial roles in the development of various diseases, such as neurological disorders and various cancers [26–28]. CircRNAs are highly stable as compared to linear RNA transcripts, and can be detected in saliva, exosomes and blood. They are therefore considered as a promising new class of biomarkers [29–31].

Although thousands of circRNAs have been identified using total RNA-seq data, the detection and characterization of circRNAs remain challenging. CircRNAs represent a small fraction of the total RNA pool and the majority of them are expressed at low levels. In addition, circRNAs share the same sequence as their linear counterparts and can only be differentiated by the sequence reads spanning the back-splice junctions. Such back-splice junction reads are rare in RNA-seq data, making up less than 1/1000 reads in a typical sequencing library. A plethora of bioinformatics approaches have been developed to predict circRNAs and measure their expression from RNA-seq data [20, 32, 33], yet the efficiency of circRNA calls and reproducibility across different approaches remain relatively low. RNAse R treatment, a process that digests all linear RNAs but preserves circRNAs, followed by PCR or RNA-seq has been instrumental for validating circRNAs identified from total RNA-seq. However, this strategy hampers the ability to quantify the abundance of circRNAs in relation to their cognate linear RNA. Finally, due to their relative short sequence and the sequence similarity with their cognate RNA, it has been challenging to define the subcellular localization of circRNAs using available in situ methods such as FISH. Defining subcellular localization of circRNAs would provide novel insights to their function. For instance, if cirRNAs are enriched in a particular subcellular compartment different from their cognate RNA, the spatial discrepancy between the two molecules could indicate independent functions and regulatory pathways.

Although circRNAs are currently under intensive investigation, there is a lack of approaches to study their function, including high- throughput methods to quantify their abundance in relation to the expression of their cognate linear mRNA. Also there is a lack of methods to define the spatio-temporal expression of circRNAs across tissues, within tissues, and at the subcellular level. To address these methodological issues, we developed a robust target-enrichment sequencing approach to provide a measure the abundance of circRNA expression their cognate linear transcripts across many samples. Furthermore, we adapted the padlock probes to visualize circRNA and to determine their subcellular localization in situ, in cells and tissues.

## Results

In this study, we developed an efficient and cost effective sequencing strategy to accurately measure circRNA expression from various input material and in a high throughput fashion. This strategy, here called the circRNA AmpliSeq panel, is based on target enrichment of circRNAs and their cognate linear isoforms using the Ion AmpliSeq™ target-enrichment sequencing panel (Figure 1A). We also report improvements for the padlock method combined with in situ sequencing to define the saptio-temporal expression of circRNAs at the subcellular level (Figure 1B).

**Figure 1:**
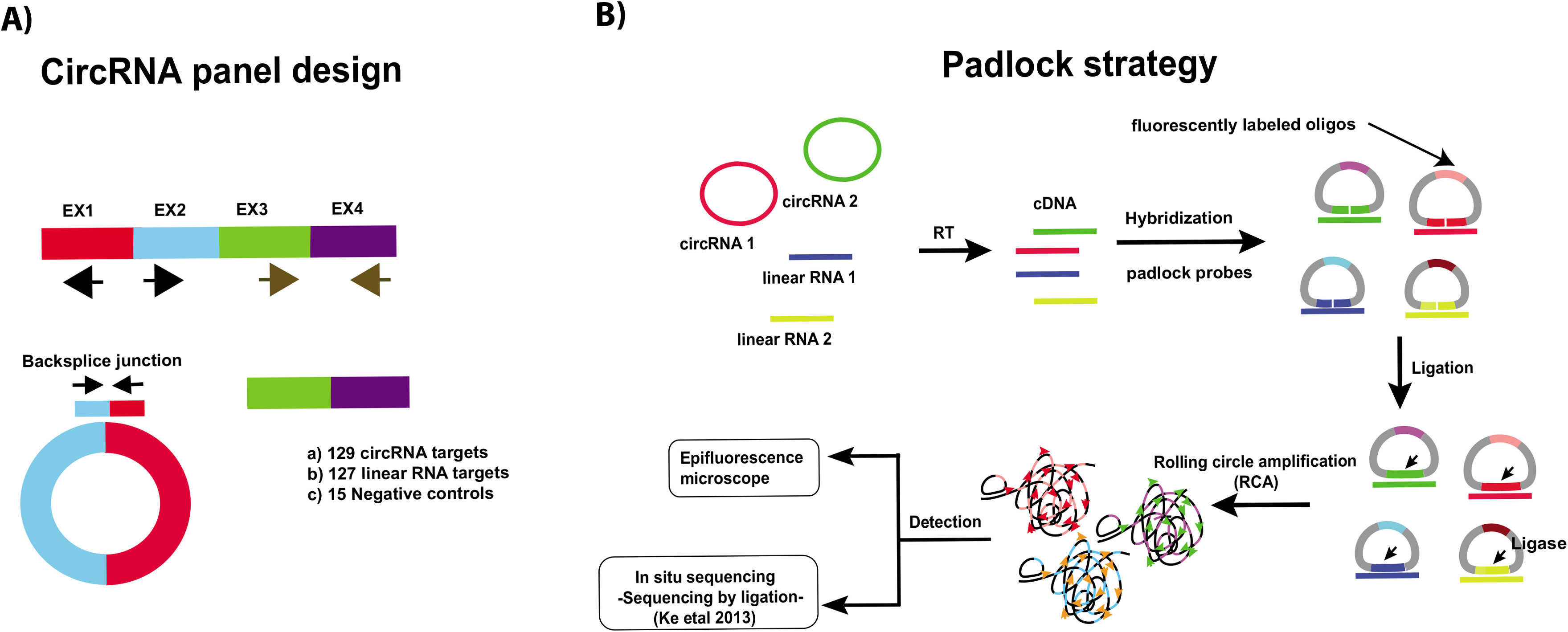
A) A schematic illustration for the design of the primers used to enrich for circRNA and their corresponding mRNA in the AmpliSeq panel. Target specific primers for sequencing-library preparation included: 129 circRNAs (primers facing outwards), 127 linear RNAs (primers facing inwards) and 15 negative controls (primer facing outwards in regions where no circRNAs are annotated. CircRNA targets were selected based on human brain RNA-seq data. B) A schematic illustration for the detection of circRNA using padlock probes and RCA. RNA is first reverse transcribed followed by hybridization of the padlock probes. Upon perfect hybridization to the target sequence, the 5’ and 3’ of the padlock probe come in proximity and circularized by target-dependent ligation. Circularized padlock probes can be amplified by RCA in situ and visualized using fluorescence microscopy or screened using in situ sequencing.

### CircRNA AmpliSeq panel design and validation

In order to identify targets to include in the AmpliSeq panel enriching for selected circular and linear RNAs, we first performed standard RNA-seq on total RNA extracted from human fetal frontal cortex, fetal liver, and fetal heart. To identify circRNAs expressed in these tissues, we then called sequence reads mapping to back-splice junctions from known exon junctions using the circRNA_finder pipeline described in [9]. For initial testing, we focused on brain tissues, as circRNAs are highly abundant and diverse in brain compared to other tissues. From the circRNAs identified in the frontal cortex, we selected 129 circRNAs expressed at high levels as well as 127 linear RNA targets from the same genes for the design of the circRNA AmpliSeq panel. Additionally, we included 15 negative control targets, i.e. back-splice junctions with no support from the RNA-seq data and with no circRNA reported in literature (see Supplementary materials for the targets and their primer sequences). The target enrichment approach for the circRNA and linear RNA is described in Figure 1A. To test if our targeted approach successfully captures circRNAs, we performed a pilot sequencing experiment using the circRNA panel on total RNA extracted from human fetal and adult frontal cortex tissues. We also used the panel to assay technical replicates of total RNA extracted from the human neuroblastoma SHSY-5Y cell line.

The sequencing generated approximately 8 million reads per sample, with 97% of the reads mapping to the panel targets. Based on sequencing read counts for all targets in the panel, we found high correlation in circRNA and linear RNA expression between technical replicates (R^2^=0.963), highlighting the reproducibility of the circRNA AmpliSeq panel (Figure 2A). To evaluate the efficiency of the circRNA panel as compared with circRNA detection from total RNA-seq data, we compared the number of sequence reads for all individual targets in the circRNA panel to the number of reads mapping to the same targets in the standard total RNA-seq data from the same samples. For this analysis, we aligned the RNA-seq reads to the exact same target sequences as defined by the AmpliSeq panel and thereby obtained an RNA-seq count for each target. Then, we directly compared the resulted RNA-seq counts to the corresponding counts obtained from the circRNA panel (Figure 2B). Our results show that the target enrichment leads to a read count several orders of magnitude higher in the circRNA panel compared with RNA-seq (Figure 2C and supplementary Figure S1), but the results of the two approaches are still correlated with R^2^ values of 0.332 ranging between 0.114 and 0.495.

**Figure 2:**
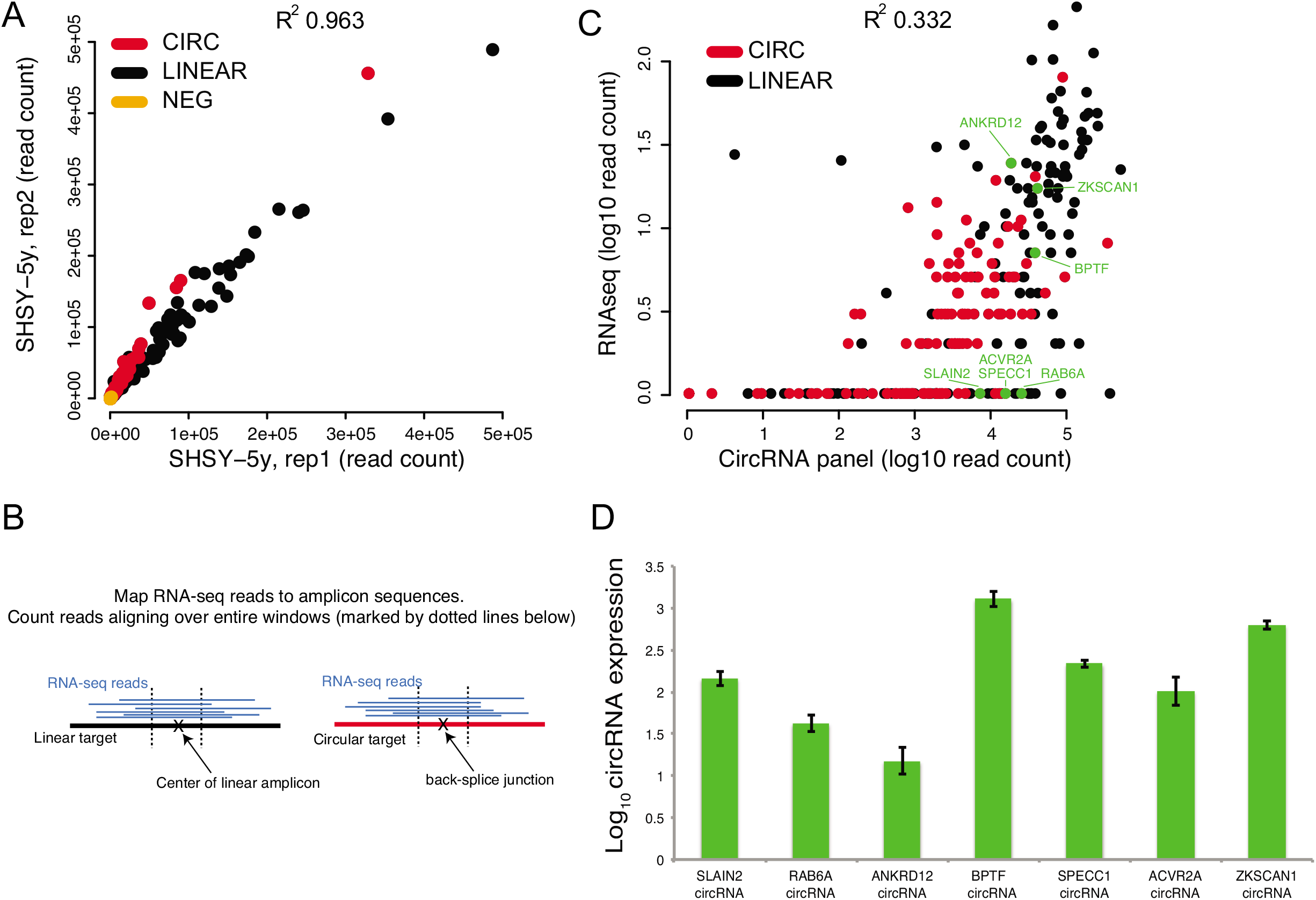
A) Correlation of expression levels (raw read counts) obtained from the circRNA AmpliSeq panel, for two replicate RNA samples from the SHSY-5Y cell line. The black dots correspond to linear RNA targets, red dots correspond to circular RNA targets, and yellow dots correspond to negative control targets. B) To make the RNA sequencing data comparable to results obtained from the circRNA panel, we aligned the RNA-reads to the reference sequences obtained from the AmpliSeq panel design and required at least 50 bases to be mapped over the middle of each amplicon in order to call a match (indicated by the dotted lines). This implies that the RNA-seq read covers at least 25bp at each side of the back-splice junction for circRNAs, and 50 consecutive bases for linear RNAs. C) Quantification of circRNA expression levels from total RNA seq (y-axis) and the AmpliSeq circRNA panel (x-axis) in SHSY-5Y RNA. The axis scales are logarithmic and the value 1 was added to each target, implying that non-expressed have a value of 0 in the plot. Each dot represents a target amplicon in the circRNA panel, either a circular RNA (red) or a linear RNA (black). D) Validation of circRNA expression. The expression level of circRNAs from SLAIN2, RAB6A, *SPECC1, ACVR2A, ZKSCAN1*, *ANKRD12 and BPTF* (represented by the green dots highlighted in 2B. In addition we included a Neg control (negative control target from the circRNA panel, as in (A)) was measured in SHSY-5Y cells using qrtPCR. All expression values were first normalized to the level of B-actin and then circRNA expression was calculated as a fold of the expression levels from the Neg control using the ΔΔCt method. Expression levels represent mean values of the technical replicates and error bars are ±SD. All expression levels are presented as log10.

The targeted enrichment and subsequent higher read coverage provides an increased sensitivity for detecting circRNAs compared to RNA-seq. Since circRNAs are generally expressed at low levels, this increased sensitivity provides a major advantage for the detection and quantification of circRNA expression without the need to perform RNA-seq with higher depth or RNase R treatments prior library preparation. Our results also indicate that several circRNAs that were not detected using standard RNA-seq are readily captured using the panel, and these may explain the relatively low correlation values between the two methods. To validate these findings using an independent method, we selected four circRNA targets (*SLAIN2*, *RAB6A, SPECC1 and ACVR2A*) with no coverage in total RNA-seq while detected at high levels in the circRNA panel (Figure 2C). In addition, we included two targets (*ZKSCAN1*, *ANKRD12 and BPTF*), which had coverage in both total RNA-seq and in the circRNA panel. We used qrtPCR to measure their actual expression levels in the original RNA sample, and found the qrtPCR results to be in agreement and correlate better with the circRNA panel data compared to RNA-seq (Figure 2D).

### CircRNA expression in human tissues

To explore circRNA expression in a large range of human tissues, we applied the circRNA panel to RNA extracted from a set of human adult and fetal tissues including frontal lobe, liver, heart, placenta and blood (n=19, Supplementary materials). To obtain an overview of circRNA expression patterns in the different tissues, we performed hierarchical clustering of the expression counts obtained from the circRNA panel for all the samples (Figure 3A). In agreement with previously published data [8, 13], we find that circRNAs are expressed in all tissues and that their expression patterns are distinguishable between the different tissues. Samples cluster by tissue type, with the exception for liver where adult and fetal samples do not cluster together. We also used the circRNA panel to measure ratio between circRNA and their cognate linear RNA and found that circRNA/linear RNA ratios vary between samples. For some genes and in some tissue types, circRNAs were expressed at higher levels than their corresponding linear RNA. In Figure 3B, we show two genes, *SETD3* and *RAB6A,* where both the absolute expression of circRNA and the ratio between circRNAs and their linear isoforms differ between the tissues. The complete table containing expression counts for linear and circular RNAs in all samples is available as a Supplementary Data file.

**Figure 3:**
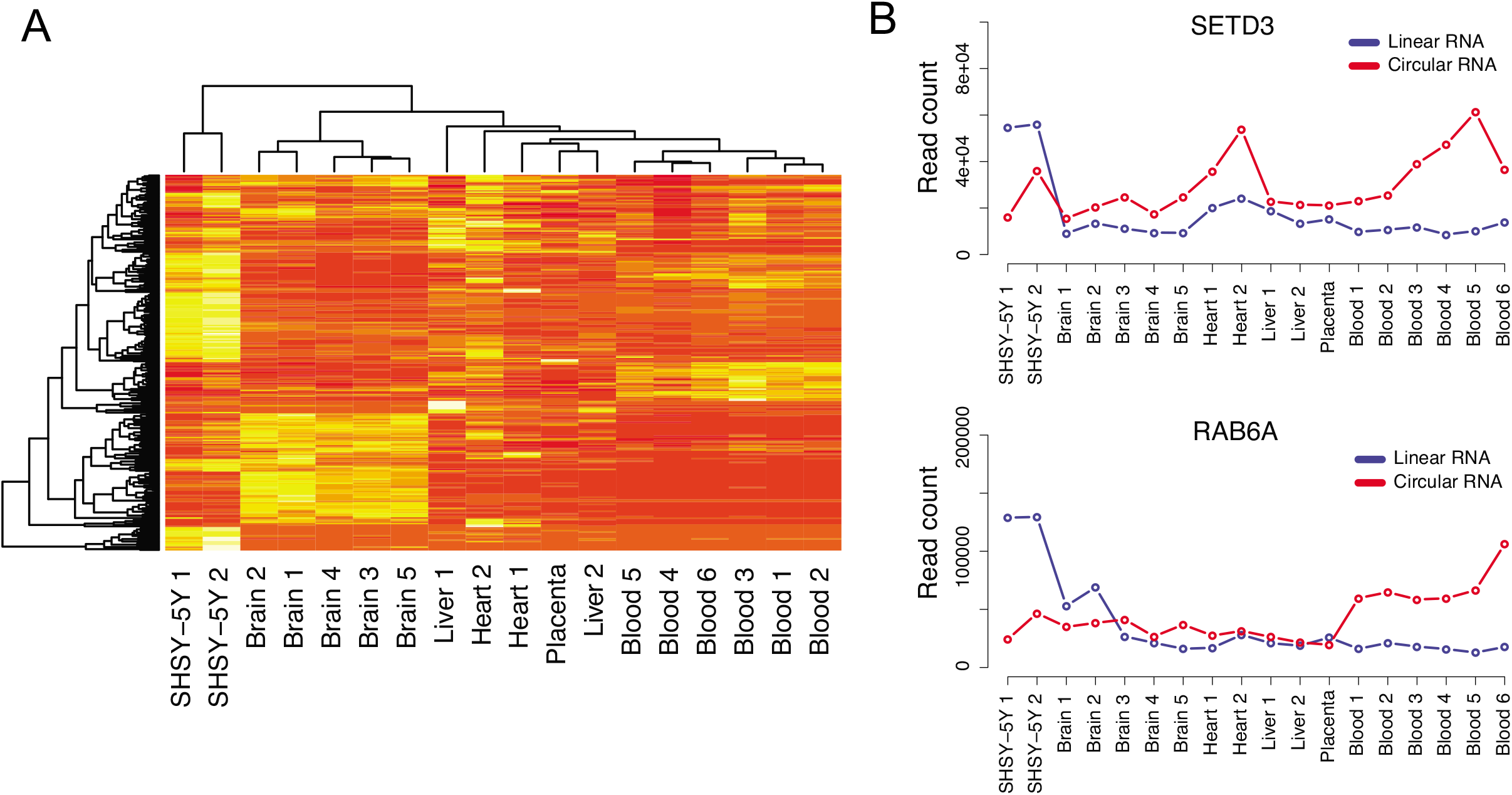
A) Hierarchical clustering different RNA samples, based on raw read counts obtained from the AmpliSeq circRNA panel. Data from all targets (circular, linear and negative control) was used for the clustering analysis. B) Read counts for the circular and linear isoforms of the genes *SETD3* and *RAB6A* in the SHSY-5Y cell line, blood and several tissue samples.

### CircRNA Expressions in FFPE and serum samples

We then tested if we are able to enrich for and detect circRNA from specimens with low RNA quality such as formalin-fixed paraffin embedded (FFPE) tissues or low RNA content such as in serum. Total RNA was first extracted from 6 FFPE tissues and 1 serum sample (Supplementary Data file) and then sequenced using the circRNA panel. In this experiment, we also performed RNA sequencing using the circRNA panel following RNAse R treatment from one of the adult frontal cortex (sample 5) as a control for the panel specificity. The relative circRNA expression (circRNA reads/(LinOut reads+circRNA reads)) in each sample is shown in Figure 4A. As expected, the RNAse R treatment resulted in the loss of the sequencing signal over the linear RNA targets without affecting the sequence coverage over the circRNA targets. This highlights the ability of the panel to enrich for circRNAs with high specificity. The results from FFPE samples showed substantial reduction of the number of circRNAs detected, as well as reduced expression relative to their linear counterparts, when compared with results from all other samples. We speculate that the reduction of the number of circRNAs detected in the FFPE samples is caused by the strong crosslinking and fragmentation caused by the tissue fixation process. More adjustment and optimization for the RNA extraction from FFPE could improve circRNA detection.

**Figure 4.**
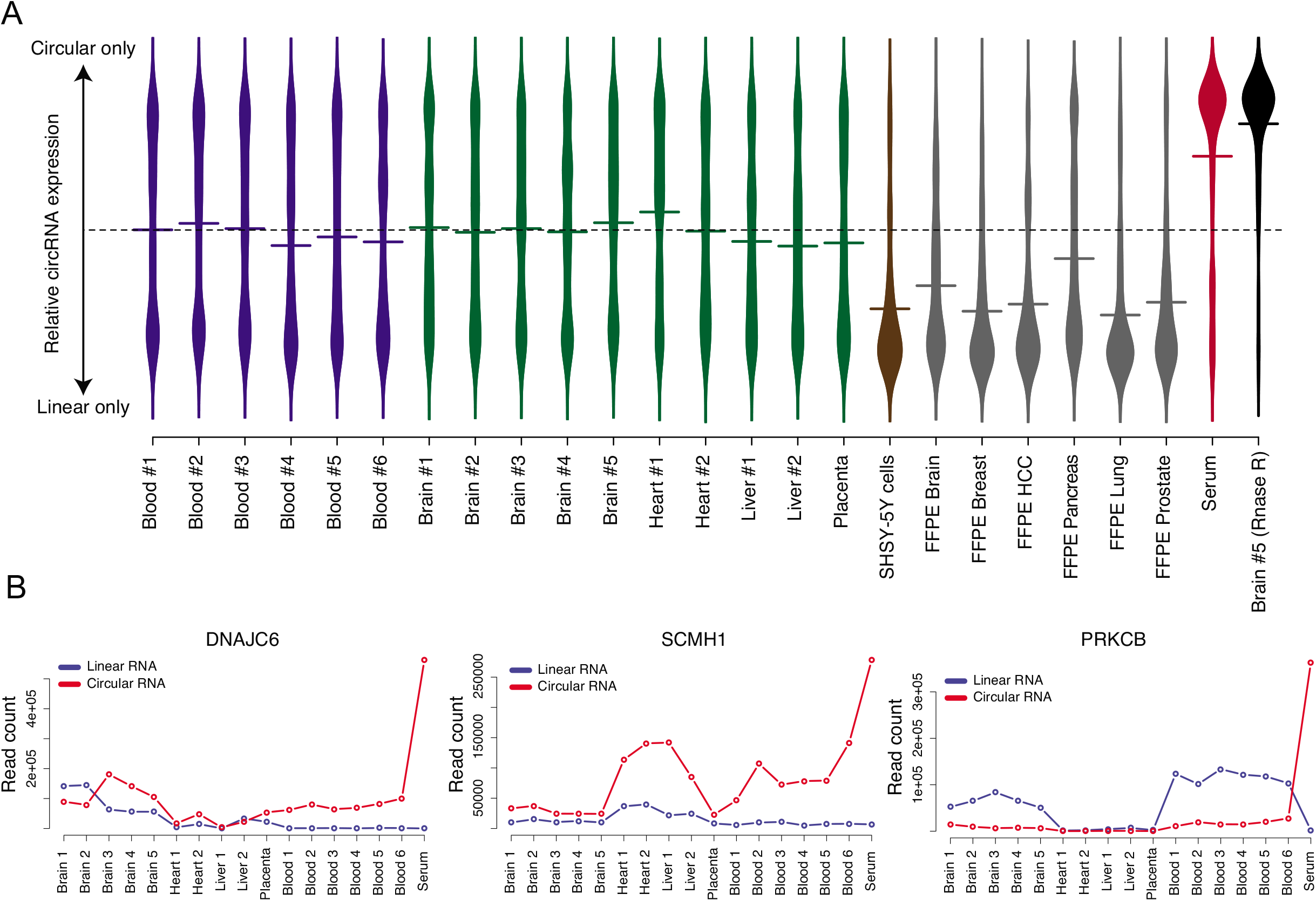
A) Violin plot showing the relative circRNA expression for all genes in the AmpliSeq panel (i.e. ratio circRNA/(circRNA+linearRNA). The average circRNA expression values are represented by horizontal lines. The dashed line represents the value where circRNA and linear RNA are equally expressed. The RNAse R treated sample was used as a control to evaluate the specificity of the circRNA panel B) Read counts for the circular and linear isoforms of the genes *DNAJC6*, *SCMH1* and *PRKCB* in tissue samples, blood and serum. All three genes show a dramatic increase of relative circular RNA expression in serum, indicating that the linear isoforms of *DNAJC6*, *SCMH1* and *PRKCB* are virtually absent from that sample.

Strikingly, we find several circRNAs to be highly expressed in the serum sample. Three examples of such circRNA are shown in Figure 4B. Expression of *DNAJC6* is of particular interest, since only the circRNA isoform is expressed in blood and serum*. DNAJC6* encodes the neuronal protein Auxilin, and it is known have a role in clathrin-mediated endocytosis and to be implicated in the pathogenesis of Parkinson disease and intellectual disability [34, 35]. Previous data from The Genotype-Tissue Expression (GTEx) project shows that *DNAJC6* is exclusively expressed in the brain [36, 37]. The detection of *DNAJC6* circRNA in the blood could indicate a role for this circRNA independent from the function of its corresponding mRNA isoform. Moreover, the presence of circRNA in serum reflects the high stability of circRNA as compared to linear RNA transcripts, which indicate the possibility of using circRNA as potential biomarkers for a particular biological or pathological condition.

### CircRNA detection using padlock probes and RCA

In order to validate our findings from the circRNA panel and to evaluate the possibility of visualizing circRNA in situ, we adapted a protocol for padlock probes in combination with rolling circle amplification (RCA) [38]. Initially, we tested this method in human fixed cells. We performed multiplex detection of three circRNAs (*CDRas1*, *HIPK3* and *MAN1A2*) using padlock probes targeting sequences surrounding the back-splice junctions followed by RCA (Figure 1B). These targets were selected due to their high expression as seen in our sequencing data, and all of them were successfully detected in SHSY-5Y cell line (Figure 5A). Although, the detection efficiency of the padlock in the cell line is less sensitive than circRNA panel, the relative abundance of the RCA signal for the circRNA targets correlated well with the expression pattern obtained from the circRNA panel sequencing data (Figure 5B). To test the method’s ability for dual detection of circRNAs and their corresponding linear RNA, we designed additional padlock probes for the *HIPK3* and *MAN1A2* linear RNA transcripts. Both the circRNA and linear RNA were simultaneously detected for both genes (Figure 5C). In accordance with our sequencing data, the padlock experiment showed higher expression for circRNAs from both genes as compared to their corresponding linear isoforms.

**Figure 5:**
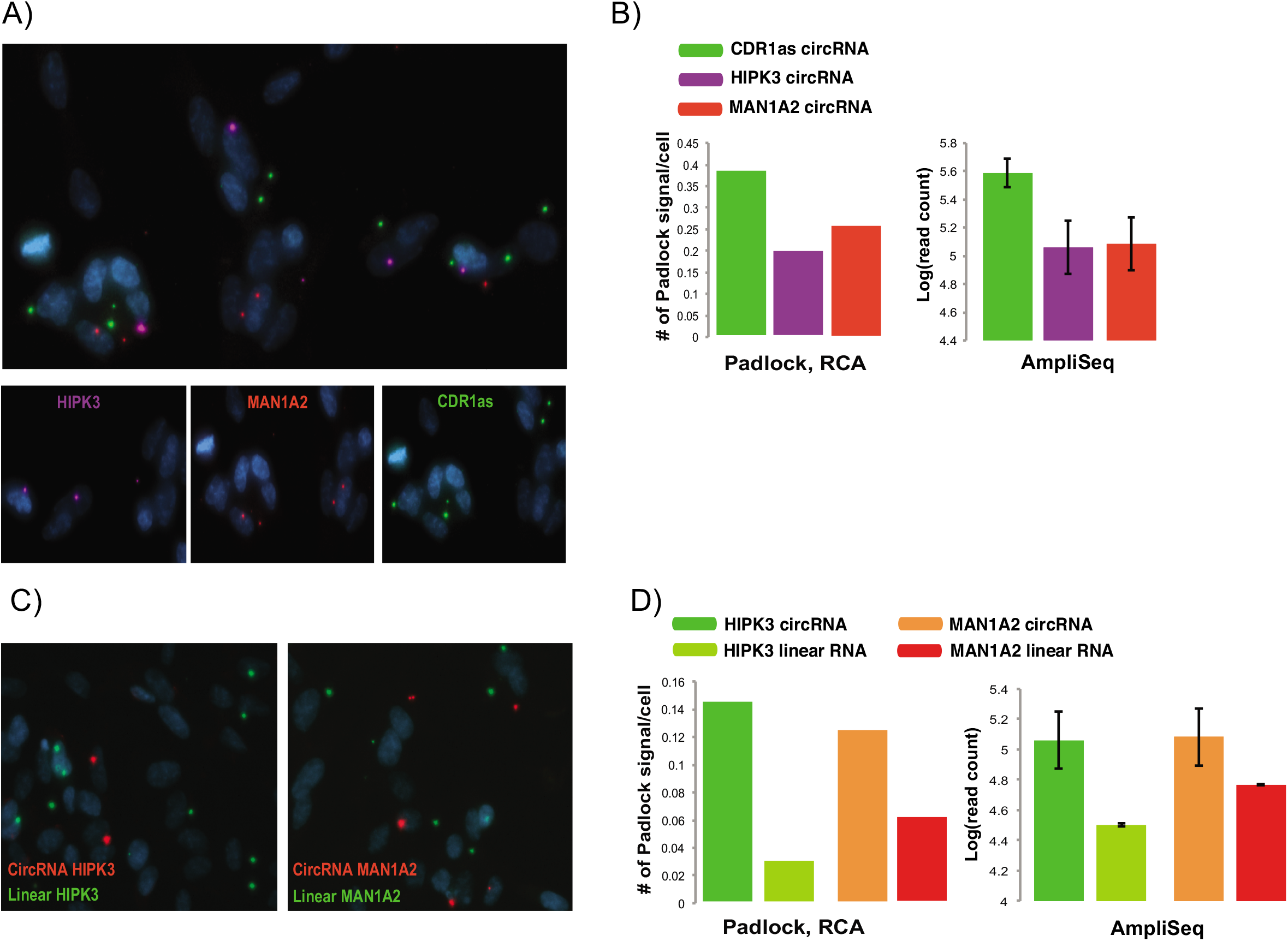
Detection circRNA and linear RNA in SHSY-5Y cell lines using padlock probes and rolling circle amplification RCA A) Multiplex detection of three circRNAs from *HIPK3*, *MAN1A2* and *CDR1as* using padlock probes complementary to the backsplice junction (Supplementary materials). RCPs from each of the circRNAs is represented with different dyes, *HIPK3;* purple, *MAN1A2*; red and, *CDR1as*; green. Top; is a merged signal from the three circRNA. Bottom; shows the signal from each of the circRNAs. B) A comparison between the padlock and RCA signal intensity and circRNAs expression from circRNA panel. Left panel shows the average RCPs/cell for circRNAs from *HIPK3, MAN1A2 and, CDR1as*, while the right panel shows log10 read count identified using circRNA panel for the same circRNAs. Similar expression patterns were observed in both strategies highlighting the reproducibility of our data. C) Dual detection of circRNA and linear RNAs form *MAN1A2* and *HIPK3* in SHSY-5Y cells. RCPs from circRNA and linear RNA are labeled in red and green fluorescence respectively. D) A comparison between the padlock and RCA signal intensity and circRNAs expression from circRNA panel. Left panel shows the average RCPs/cell for circRNA and linear RNA from *HIPK3* and *MAN1A2*, while the right panel shows log10 read count identified using circRNA panel for the same circRNAs.

### Subcellular localization of circRNA in the brain

While it is believed that the majority of circRNAs are cytoplasmic, the detection of circRNAs in the nucleus is also reported [18]. Since current reports suggest that circRNAs could display functions independent from the function of their host gene, we motivate that it is interesting to investigate if some circRNAs show different localization than their cognate linear RNA molecules. Therefore, we extended the use of the padlock approach and rolling circle amplification (RCA) to study the subcellular localization of circRNA in vivo.

To define the cellular localization of circRNAs, we applied an in situ sequencing approach, which in principle allows the detection 273 targets in one experiment, via the application of padlock probes and RCA [39]. We targeted highly expressed circRNAs detected in our brain RNA sequencing data and investigated their subcellular localization in a fresh/frozen brain tissue section. The tissue section is from the same tissue used to isolate RNA for the circRNA panel (sample 5, see supplementary materials). In this experiment, we designed padlock probes targeting 20 bp sequences overlapping and surrounding the backsplice junction of 8 circRNA targets (see supplementary materials for padlock probes sequences) (Figure 1A). We also targeted their corresponding mRNA by targeting exons downstream of the backsplice junction. As a control for the cellular localization experiment, we included padlocks targeting the *Malat1* lncRNA, which is predominantly expressed in the nucleus [40]. Following RCA of the specifically-bound padlock probes, we subjected the RCA products (RCPs) from each target to sequencing-by-ligation [39]. Each individual padlock probe harbored a unique barcode (4 nucleotides) to distinguish different targets [39] (Figure 6A). We first plotted the number of sequence reads from the in situ sequencing for each target and then assigned them to their subcellular localization. As expected, in situ sequencing results indicated that the linear *Malat1* transcript is predominantly expressed in the nucleus (Figure 6B, and Supplementary figure S2), while the circRNAs *CDR1as*, *HIPK3* and *MAN1A2* were enriched in the cytoplasm, as previously reported [10, 41, 42]. Of the eight circRNA targets tested, six were found to be enriched in the cytoplasm (Figure 6C, and Supplementary figure S2). The circRNA target *SNAP47* showed equal distribution between nuclear and cytosolic compartments, while *MGAT5* showed predominantly nuclear localization.

**Figure 6:**
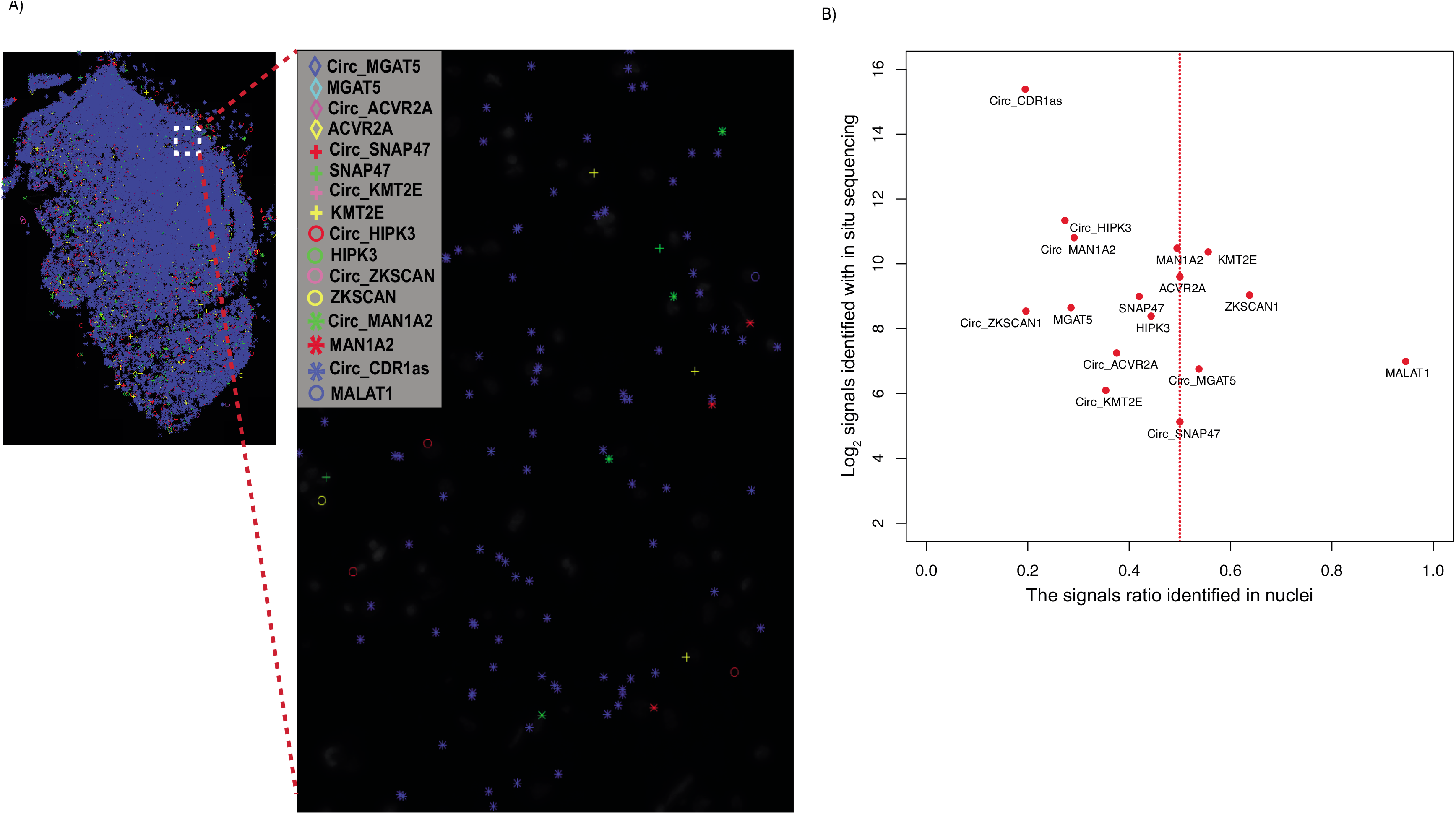
Multiplex detection of circRNAs and linear RNAs with padlock probes in combination with in situ sequencing on human brain tissue. A) Padlock probes, RCA, and in situ sequencing were performed to detect the cellular localization of circRNAs and linear RNA in a tissue section from fresh/frozen human brain (sample 5). CircRNAs and their corresponding linear RNA from 7 genes were screened (*MGAT5, ACVR2A, SNAP47, KMT2E, HIPK3, ZKSCAN*, and *MAN1A2*). In addition, circRNA from *CDR1as* and linear RNA from *MALAT1* (known to be exclusively expressed in the nucleus, and used as a control for the localization experiment) were also included. Padlock probe sequences for each of the targets are listed in supplementary materials. In situ sequencing was performed to identify the 4-bp specific sequence barcodes in RCA products (RCPs) of each of the targets as shown in [39]. All targets were plotted on the DAPI image of brain tissue section after in situ sequencing. Localization of each of the detected targets is shown as a symbol, left panel. Each symbol represents a barcode sequence that corresponds to a specific target. The nuclei are shown in grey. Scale bar, 100μm. Zoom in region corresponds to the region in white dash-line square. B) Abundance and subcellular localization of circRNA and linear RNAs. The ratios of signals in nuclei for all targets were plotted against their barcode counts in the brain sample. Similar to the circRNA panel data, *CDR1as* was the most abundant circRNAs in the in situ sequencing experiment. Also, in agreement with previous data, *MALAT1* was highly expressed in the nucleus in the in situ sequencing experiment.

## Discussion

Although there is an increasing interest in unraveling the function of circRNAs and understanding their role in human disease, there is a lack of efficient and reliable tools to study their basic characteristics such as their expression levels-especially in relation to their cognate linear mRNA, and their cellular localization. In this study, we developed two approaches to allow an improved understanding of the nature and function of circRNAs. In the first approach, we aimed to develop an efficient and cost effective tool to characterize the expression patterns of circRNAs in different tissues and cells. To achieve this, we designed a pilot target-enrichment sequencing panel using the Ion AmpliSeq^TM^ technology to enrich for circRNAs known to be expressed in human brain and their corresponding mRNA transcripts. We this panel, we produced an average of 8 millions reads/sample which resulted in a coverage over circRNAs several orders of magnitude higher than RNA-seq with 40 million reads/sample. In the second approach, we adapted the padlock technology in combination with RCA and in situ sequencing to detect and identify the spatiotemporal expression of circRNAs at the subcellular level in cells and fresh/frozen tissue sections.

The ability to quantify the abundance of circRNAs in various tissues and to identify how their expression is correlated with their corresponding mRNA is crucial to understand the nature and the function of circRNAs. Using the circRNA panel, we successfully enrich for the circRNAs and their corresponding mRNAs from various input materials. We find the circRNA panel is more efficient at detecting circRNAs than total RNA-seq. The target enrichment approach leads to significantly higher sequence coverage over circRNA and mRNA targets as compared to RNA-seq. This enabled the detection of circRNAs with very low (or no) read coverage in RNA-seq data, leading to improved and more sensitive and comprehensive estimation of circRNA expression, especially for circRNA expressed at low levels or in specimens that contain a low amount of RNA. The target enrichment allowed us to screen multiple samples (n=19) using one P1 Ion Proton sequencing chip, producing approximately 80 million reads/P1 chip. On the other hand, the total RNA-seq for similar samples in our screening required approximately 40 million reads/sample (2 samples/chip), highlighting the cost effectiveness of circRNA panel over total RNA-seq. It is also important to note that the circRNA panel designed for this study is a proof-of-concept pilot panel, created only to show the feasibility of high throughput screening. It is possible to scale this up to detect thousands of circRNA and linear RNA targets using a single panel.

Using the circRNA panel to analyze circRNA and mRNA expression in various tissues we found, in accordance with previous reports, that circRNAs show distinct expression patterns between different tissues. Moreover, the circRNA to mRNA expression ratio exhibited different patterns among the different samples. Detection of circRNA in FFPE samples was particularly inefficient. Although we could still detect a small number of highly expressed circRNAs in FFPE samples (Supplementary materials), there was a clear reduction of circRNA as compared to their linear counterparts. These results indicate that circRNAs are not present in extracted total RNA from FFPE, either due to degradation or the general challenges associated with RNA purification from FFPE samples.

We found several examples where the circRNA isoform is highly expressed, while the linear mRNA isoform is not expressed or expressed at very low levels. One example is *DNAJC6*, a neuronal protein where the mRNA, in our data and in previous studies, is expressed exclusively in the brain [36], while we show for the first time that its circRNA isoform is expressed at high levels in blood, and also detectable in serum. These results support a possible role of circRNAs as stable tissue-specific biomarkers accessible in serum. The findings also highlight the efficiency and feasibility of our circRNA panel to further evaluate and investigate the role of circRNA as biomarkers for diagnosis and treatment.

Prior to this study, defining the cellular localization of circRNA and their cognate linear mRNA in the same experiment experiment using common approaches, such as FISH, was hampered due to the sequence similarity between circRNA and their corresponding mRNA and because circRNAs are typically composed of short sequences. Similar to other non-coding RNAs, defining the subcellular localization of circRNA could provide valuable insights into their function. For instance, cytoplasmic circRNAs are suggested to regulate gene expression through interfering with mirRNA pathways. While nuclear circRNAs, which mostly represent intron-containing circRNAs, are suggested to interact with the splicing machinery and promote transcription [18]. In this study, we demonstrate that the padlock approach followed by RCA can be used to visualize circRNA and their corresponding mRNA in cells and tissue sections. The abundance of circRNA obtained from the padlock experiment from the SHSY-5Y cell line correlated well with the results from the circRNA panel. However, we believe that further optimization is possible to increase the sensitivity for detecting circRNAs expressed at moderate and low levels. When combined with in situ sequencing (sequencing-by-ligation), this approach allowed multiplex detection of circRNA and their corresponding mRNA in human brain at the subcellular level. These results highlight the possibility of using our in situ sequencing strategy to define the subcellular localization of circRNAs and how they co-localize with their corresponding mRNA isoforms. To our knowledge, this strategy is the first of it is kind to allow a multiplex detection and subcellular localization data of circRNAs and their corresponding mRNAs in situ at the subcellular level. In addition, it offers an opportunity to resolve the spatio-temporal expression of circRNA in complex tissues such as the brain.

Both the circRNA panel and the circRNA padlock visualization approaches presented here will pave the way for an improved hypothesis-driven targeted analysis of circRNA functions in cells and tissues. The enrichment of the circRNA panel allows a cost-effective approach to perform surveys of circRNA expression profiles in large sample sets. The circRNA panel setup offers the possibility to increase the number of targets and to perform custom design for the circRNAs of interest. On the other hand, the padlock strategy offers, for the first time, an experimental approach to perform multiplex detection of circRNA cellular localization in vivo. This approach can also be used to define the spatial context of circRNA expression profiles between different cell populations within the same tissue. In conclusion, our approaches represent powerful a toolbox for deep profiling and characterization of circRNA and for unlocking their roles as potential diagnostic biomarkers.

## Methods

### Sample preparation for total RNA sequencing

Total RNA from fresh frozen tissues (fetal frontal lobe, fetal liver and fetal heart) was purchased from Capital Biosciences. Total RNA from fetal tissues and SHSY-5Y cells was extracted using RiboPure^TM^ RNA purification kit (Ambion) according to the manufacturer instructions.

### Total RNA sequencing

RNA was treated with the Ribo-Zero Magnetic Gold Kit for human/mouse/Rat (Epicentre) to remove ribosomal RNA, and purified using Agencourt RNAClean XP Kit (Beckman Coulter). The rRNA depleted RNA was then treated with RNaseIII according to Ion Total RNA-Seq protocol v2 and purified with Magnetic Bead Cleanup Module (Thermo Fisher). Sequencing libraries were prepared using the Ion Total RNA-Seq Kit for the AB Library Builder™ System and quantified using the Fragment analyzer (Advance Analytical). The quantified libraries were pooled followed by emulsion PCR on the Ion OneTouch 2 system, and sequenced on the Ion Proton System.

### Discovery of circular RNAs in total RNA-seq data

To identify circular RNAs in total RNA data, a computational pipeline adapted from [9] was used. In short, reads were aligned using STAR [43], allowing for chimeric transcripts. The candidate chimeric junction reads were then filtered to only include cases where the read spanned a junction with the splice acceptor on the same chromosome and strand as the splice donor, at most 100,000 bp upstream. In the subsequent analysis only circular junctions matching GT-AG splice sites we included.

### Design of an AmpliSeq panel for circular RNA

Loci to be included in the circRNA ampliseq panel were selected using the following criteria: First, circRNAs with high expression in total RNA-seq data from brain were chosen, so that each circRNA originated from a different mRNA transcript (i.e. a backsplice site between annotated exons). We required the length of each circRNA (assuming introns are spliced out) was at least 300 nt. Controls measuring expression of linear RNA isoforms (LinOut) were placed on the same transcript 300 nt upstream from the circRNAs if possible, otherwise 300 nt downstream. As negative controls, possible back splice sites with no support from the RNA-seq data were sampled, from transcripts expressed in the brain. The only exception to this scheme was CDR1as, which was added manually to the panel and doesn’t have a control for linear mRNA, as it does not originate from a known mRNA transcript. Design of the circRNA panel was based on the hg19 genome and transcript models from the UCSC genome browser.

### Sample preparation for sequencing using the circRNA AmpliSeq panel

Adult brain samples (sample 1-5, Supplementary materials) were obtained from the Harvard Brain Tissue Resource. Total RNA was extracted from 5 adult frontal lobe samples. To obtain total RNA, tissue pieces were first fixed in OCT and two sections were used for total RNA purification. Total RNA purification for these samples and 6 human blood samples was performed using RiboPure^TM^ RNA purification kit (Ambion) according to the manufacturer instructions. Total RNA from fetal and adult liver, fetal and adult heart, and placenta were purchased from Clonetech. Total RNA from the FFPE tissues was extracted using High Pure FFPET RNA isolation kit (Roche) according to the manufacturer instructions. RNA from the serum was purified using Plasma/Serum Circulating and Exosomal RNA purification Kit (Norgen). RNase R treatment of total RNA was performed after ribosomal RNA depletion. Briefly, 2 μg RNA was added to a mixture of 1x RNase R buffer (epicentre®), 1mM MgCl_2_, and 1.0 μl (20U/μl) RNase R enzyme (epicentre®). The mix was then incubated at 37C° for 10 minutes. RNase R was then inactivated by adding 1:1 volume of H_2_O. All samples are listed in Supplementary materials.

### Sequencing using the circRNA panel

RNA was reverse transcribed to cDNA, and the acquired cDNA was amplified using the custom made Ion AmpliSeq™ circRNA panel. The sequencing libraries were prepared using the Ion AmpliSeq™ Library Kit 2.0 (Thermo Fisher). Sequencing libraries were purified using the Agencourt AMPure XP reagent (Beckman Coulter) and amplified. The amplicons were quantified using the Fragment Analyzer instrument (Advanced Analytical). Samples were then pooled, followed by emulsion PCR using the Ion Chef System, and sequenced on the Ion Proton System.

### qrtPCR validation

Starting with 1 μg of total, cytoplasmic, nuclear RNA, cDNA was synthesized using the RevertAid First strand cDNA synthesis kit (Fermentas) with random hexamers according to the manufacturer’s recommendations. 1 μl of the resulting cDNA was used for qrtPCR to measure the relative circRNA/linear RNA ratio in each sample. The qrtPCR was performed with Stratagene Mx3000P in 96-well plates. The reactions were carried out with an initial denaturation at 95°C for 10 min followed by 40 cycles of denaturation at 95°C for 15 s, primer annealing at 60°C for 30 s and extension at 72°C for 30 s. The qrtPCR contained 12.5 ng single stranded cDNA, 0.4 μM for each primer and 12.5 μl Maxima SYBR Green/ROX qPCR Master Mix (Fermentas) in 25 μl reactions. All samples were amplified in triplicate and the mean values were used to calculate the expression level of each target. The circRNA RNA expression level were determined using the ΔΔCT method. CT values were first normalized to the levels of B-actin. Then, expression values for circRNA expression after normalization were calculated as a ratio between the circRNA expression values and the expression values of the negative control. Raw data were analyzed using MxPro Mx3000P software (Stratagene).

### Preparation of cells and tissue sections for circRNA visualization

SHSY-5Y cells were cultured in 1:1 mixture of Ham’s F12 and DMEM (Gibco) medium (Gibco) with L-glutamine and phenol red, supplemented with 2 mM l- glutamine (Sigma), 10% FBS (Sigma) and 1× PEST (Sigma). All cell lines were incubated at 37 °C, 5% CO2. To prepare cell slides for the padlock experiment, cells with >80% confluence were treated with 0.25% (w/v) trypsin-EDTA (Sigma) and resuspended in culturing medium. Thereafter, cells were seeded on Superfrost Plus slides (ThermoFisher) and placed in a 150 mm × 25 mm Petri dish (Corning), and culturing medium was added to a final volume of 25 ml. Cells were then incubated for 12 h at 37 °C, 5% CO2. Cell fixation was then performed in 3.7% (w/v) paraformaldehyde (Sigma) in DEPC-treated PBS for 20 min at RT. Following the fixation, cell slides were washed twice in DEPC-treated PBS and dehydrated in an ethanol series of 70%, 85% and 100% for 5 min each. Then cell slides were kept at -80 °C until use. Tissue sections (4 μm) from human fresh-frozen frontal lobe (Sample 5, Supplementary materials) were prepared with microtome and mounted on a Superfrost Plus slides (ThermoFisher). Tissue samples were stored at -80 °C until fixation. Fixation was performed in 3.7% (w/v) paraformaldehyde (Sigma) in DEPC-treated PBS with 0.05% Tween-20 (Sigma) (DEPC-PBS-T) for 45 min at RT and two washes in DEPC-PBS-T. The tissue sections were then permeabilized in 2 mg/ml pepsin (Sigma) in 0.1 M HCl at 37 °C for 2 min. After two washings in DEPC-treated PBS and dehydration in an ethanol series, as in the cell line, tissue sections on slides were then covered with Secure-Seal hybridization chambers (Invitrogen).

### In situ reverse transcription, barcode padlock probing and rolling circle amplification (RCA)

Cells and tissue samples were first rinsed with DEPC-PBS-T. A reversed transcription mix, containing 5 μM of unmodified random decamers, 0.2 μg/μl BSA (NEB) 500 μM dNTPs (Fermentas), 20 U/μl of TranscriptMe reverse transcriptase (Gdansk), and 1 U/μl RiboLock RNase Inhibitor (Fermentas) in the TranscriptMe reaction buffer, was applied to the slides. The incubation was carried out for 3 h at 37 °C for the cell line and overnight for the tissue sections. Slides were washed twice with DEPC-PBS-T, and followed by a post-fixation step in 3.7% (w/v) paraformaldehyde in DEPC-PBS for 10 min (or 30 min for tissues) at room temperature. After post-fixation, the samples were washed twice in DEPC-PBS-T. Following the reverse transcription, RNA degradation, hybridization and ligation of padlock probe on synthesized cDNA were performed. A mix contained 1× Ampligase buffer (20 mM Tris-HCl, pH 8.3, 25 mM KCl, 10 mM MgCl_2_, 0.5 mM NAD and 0.01% Triton X-100), 100 nM of each padlock probe, 50 μM dNTPs, 0.5 U/μl Ampligase, 0.4 U/μl RNase H (Fermentas), 50 mM KCl and 20% formamide was added into the sample reaction chambers on slides. The slide was incubated at 37 °C for 30 min, and 45 °C for 45 min, and then washed twice with 1× DEPC-PBS-T. For RCA, the reaction chamber was incubated with a RCA mix (1 U/μl phi29 polymerase (Fermentas), 1× phi29 polymerase buffer, 0.25 mM dNTPs, 0.2 μg/μl BSA and 5% glycerol in DEPC-H_2_O) for 2.5 h washed twice in DEPC-PBS-T. For tissue slide, RCA was carried out overnight.

### Sequencing-by-ligation

An ethanol wash series were firstly used to remove the mounting medium on tissue sections. As previous study [39] slides were washed with DEPC-PBS-T once and treated with UNG buffer (1× UNG buffer (Fermentas), 0.2 μg/μl BSA, 0.02 U/μl UNG (Fermentas) for 30 min at 37 °C. Slides were washed twice with DEPC-PBS-T, and then washed three times with 65% formamide for 60s each. In sequencing-by-ligation chemistry, anchor primers and interrogation probes were hybridized separately before the ligation. Briefly, a hybridization mix containing 500 nM of anchor primers in 2× SSC and 20% formamide were applied to the sample and incubated at RT for 30 min, and followed by DEPC-PBS-T wash twice. A ligation mix containing each interrogation probe, 100 ng/ml of DAPI, 1 mM ATP (Fermentas), 1× T4 ligase buffer (Fermentas), and 0.1 U/μl of T4 ligase (Fermentas), was added to the samples and incubated for 30 min at RT. Following 3x DEPC-PBS-T wash, the slides were mounted in SlowFade Gold Antifade Mountant (ThermoFisher).

### Image acquisition and analysis

Images were acquired using an AxioplanII epifluorescence microscope (20× objective). Exposure times for all the experiments are listed in Supplementary materials. After imaging, the slides were prepared for the next three sequencing cycle by UNG treatment buffer as described above followed by repeating the hybridization, ligation and imaging processes. The image analysis was performed as described in [39], a stack of images at different focal depths were captured and merged to a single image, and followed by a automatically stitching in the Zeiss AxioVision software. The fully automated sequence decoding was performed similar as previous study [39]. In short, cell nuclei were separated based on shape descriptors. Definition of cell cytoplasm uses nucleus as a seed and depends on sufficient cytoplasmic autofluorescence. The image of the general stain was enhanced by a top-hat filter, and RCPs were separated by watershed segmentation. The images were aligned, and fluorescence intensity from each of the signals representing A, C, T and G were extracted. The optimal transformation between a merged image of all signals (A + C + T + G) and the general stain was determined based on intensity [44]. Analysis were performed with CellProfiler (2.1.1, 6c2d896) calling ImageJ plugins from Fiji for image registration. All intensity information was decoded using a script written in Matlab. Briefly, RCP was assigned for the base with the highest intensity for each RCP in all hybridization steps. A quality score was extracted from each base, and the quality of a transcript was defined as the lowest quality of all the bases in the transcript. The quality score ranges from 0.25 (poor quality) to 1 (good quality). The frequency of each sequence was extracted after a typical quality threshold of 0.4-0.55 was set.

## Author contribution

AZ, AA and LF conceived and designed the study. AZ performed sample preparation, extraction and PCR validation. AZ, CW and MN designed and performed in situ experiments. AA, JOW and AN performed bioinformatic analyses. MM and KB created the AmpliSeq panel. AZ, AA, CW and LF wrote the paper with help form all authors. All authors read and approved the final manuscript.

## Acknowledgments

We thank the Uppsala Genome Center for technical support with library preparation and sequencing experiments. Computational analyses were performed on resources provided by SNIC through Uppsala Multidisciplinary Center for Advanced Computational Science (UPPMAX). This work was supported in part by the Swedish Research Council Grants 2012-4530, 2016-03645, and the European Research Council ERC Starting Grant Agreement n. 282330 (to LF).

